# Impact of *O*-GlcNAcylation elevation on mitophagy and glia in the dentate gyrus

**DOI:** 10.1101/2024.09.19.613771

**Authors:** Joshua Kramer, John C Chatham, Martin E Young, Victor Darley-Usmar, Jianhua Zhang

## Abstract

*O*-GlcNAcylation is a dynamic and reversible protein post-translational modification of serine or threonine residues which modulates the activity of transcriptional and signaling pathways and controls cellular responses to metabolic and inflammatory stressors. We and others have shown that *O*-GlcNAcylation has the potential to regulate autophagy and mitophagy to play a critical role in mitochondrial quality control, but this has not been assessed *in vivo* in the brain. This is important since mitochondrial dysfunction contributes to the development of neurodegenerative disease. We used mito-QC reporter mice to assess mitophagy in diverse cells in the dentate gyrus in response to pharmacological inhibition of OGA with thiamet G which leads to elevation of protein *O*-GlcNAcylation. We demonstrate that mitophagy occurs predominantly in the GFAP positive astrocytes and is significantly decreased in response to elevated *O*-GlcNAcylation. Furthermore, with increased *O*-GlcNAcylation, the levels of astrocyte makers GFAP and S100B, and microglial cell marker IBA1 were decreased in the dentate gyrus, while the levels of microglial cell marker TMEM119 were increased, indicating significant changes in glia homeostasis. These results provide strong evidence of the regulation of mitophagy and glia signatures by the *O*-GlcNAc pathway.

## Introduction

Mitochondria play a crucial role in cellular health and function, especially in the brain. Mitochondrial dysfunction has been implicated in neurodegenerative diseases and proposed as a cascading pathogenic factor in Alzheimer’s disease (AD) ^1-3^. In these conditions, compromised mitochondrial function can lead to energy deficits, increased production of reactive oxygen species, and a disruption in calcium homeostasis, all of which contribute to neuronal degeneration^4^. Since mitochondrial quality control depends on the lysosomal mediated process of degrading damaged mitochondria known as mitophagy, its modulation has been explored as a factor in the development of neurodegenerative diseases ^5^. In addition to neurons, recent studies have identified key roles for mitochondrial function and mitophagy in glia ^6-8^. Microglia, the resident immune cells of the central nervous system, and astrocytes, the most abundant glial cells, provide nutrient support to neurons, clear cell debris by phagocytosis, and participate in the inflammatory response. Consequently, mitochondrial dysfunction in microglia and astrocytes has been proposed to play a significant role in the pathogenesis of neurodegenerative diseases ^6,7,9-16^. How mitophagy is regulated in these cells is not fully understood.

Cellular regulation of mitophagy is also impacted by post-translation modifications, including protein *O*-GlcNAcylation, although the specific mechanisms and extent of its impact on mitophagy remain unclear^17-20^. *O*-GlcNAcylation occurs when *O*-GlcNAc is adducted to serine or threonine residues of nuclear, cytoplasmic, mitochondrial, or transmembrane proteins by *O*-GlcNAc transferase (OGT) ^21^. The modification is removed by *O*-GlcNAcase (OGA) with the net level of protein *O*-GlcNAcylation controlled by the dynamic interaction of OGT and OGA ^22,23^. We and others have shown that *O*-GlcNAcylation plays a role in virtually all cells, controlling the localization, stability, regulation, and function of proteins and signaling processes in both physiological and disease states ^21,24^. In the brain, OGT and OGA are highly expressed ^21^, and *O*-GlcNAcylation has been shown to regulate various aspects of neural function, including cell survival, autophagy ^24^, and synaptic strength ^4^. Aberrant levels of *O*-GlcNAcylation have been associated with diseases such as diabetes, cardiovascular disease, AD, and cancer ^21,24,25^.

To investigate the impact of *O*-GlcNAcylation on neuronal function, a potent and highly selective inhibitor of OGA thiamet G (TG), was used and shown to have profound effects on neurodegenerative pathways and cognitive functions ^26^. For example, it has been demonstrated that TG can prevent cognitive decline and amyloid plaque formation in tau/APP mutant mice, highlighting its potential in modulating pathways involved in neurodegeneration, although it is unclear whether this can translate to AD ^22,23,27^. Interestingly, our recent studies have shown that the effects of TG are sex-dependent and modify the relationships between gene expression, behavior, bioenergetics, and autophagy ^24,25^. These studies collectively suggest that TG, by elevating *O*-GlcNAcylation, impacts key cellular processes including mitochondrial function and neuronal health. Given that mitophagy is crucial for mitochondrial quality control and the maintenance of cellular homeostasis, we hypothesized that TG would modulate mitophagy in the brain.

In the current study we investigate the effects of elevation of protein *O*-GlcNAcylation by TG on mitophagy and glial homeostasis in the dentate gyrus (DG) of both male and female mice. The DG was selected since it is an integral part of the hippocampus, plays a central role in cognitive functions, and is implicated in neurodegenerative processes ^28^. For measurement of mitophagy, we used male and female mito-QC mice, which allow detection of mitochondria that are engulfed by autophagosomes and delivered to the lysosomes for degradation ^29^. Building on our previous work demonstrating the use of mito-QC mice in visualizing cardiac mitochondrial dynamics in vivo ^30^, the current study extends the application of this model to the DG. We provide compelling evidence that the majority of the mitophagy events in the DG occur in GFAP positive astrocytes. Within 3 hours of *O*-GlcNAc elevation, mitophagy is significantly decreased. This decreased mitophagy is observed concurrent with a decrease in GFAP, S100B, and IBA1 levels, and an increase in TMEM119 level. Taken together, these changes suggest a substantial remodeling of the glial milieu in the DG, underscoring the rapid and dynamic response of glial cells to alterations in *O*-GlcNAcylation.

## Materials/Methods

### Animals

Mito-QC mice were obtained from Dr. Ian Ganley (University of Dundee, C57BL/6) ^29^. In these mice, mCherry-GFP is fused to a mitochondrial targeting sequence from FIS1 protein. In functioning mitochondria, both mCherry and GFP proteins fluoresce. When mitophagy is initiated and a portion of the mitochondria is engulfed by the lysosome, the lowered pH environment results in the quenching of GFP fluorescence, leaving only the mCherry signal.

A total of 20 male and female mito-QC mice, aged 3 months, were provided ad libitum food and water, housed in type II polycarbonate static microisolation caging in groups of 5. The animals were sorted by sex, and then block randomization (n=5 per injection group) was used to determine which injection solution each mouse would receive. Mice were then injected i.p. with either saline or the *O*-GlcNAcase inhibitor TG (50 mg/kg), at 7 am. Six- and 26-month-old C57BL/6 (RRID:IMSR_JAX:000664) male mice were also injected with saline or TG at 7 am (n=4 each). Animals were anaesthetized by 3% isoflurane 3 h later and euthanized by saline intracardial perfusion and verified by heartbeat, using the study design shown in **Fig. 1A**. No blinding was performed in this study and no statistical methods were used to determine sample size. The study was not pre-registered. No exclusion criteria were pre-determined, and no animals were excluded from the study. All animal experiments were approved by the University of Alabama at Birmingham IACUC.

**Fig. 1:**
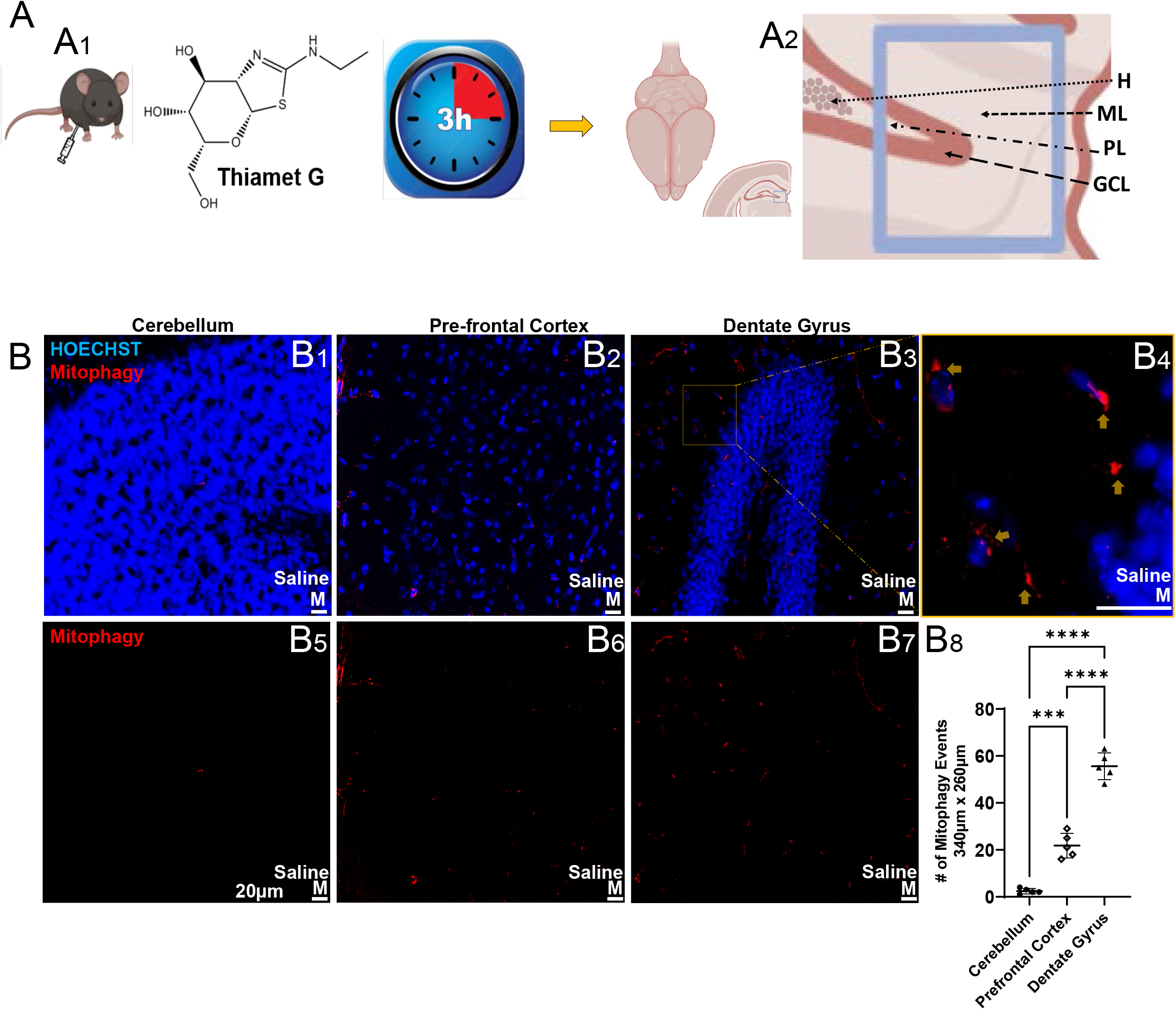
Detection of mitophagy. **A**) Experimental design. Acute thiamet G (TG) (50 mg/kg) or saline was injected in 20 male and female 3-month-old mice at 7 am. Animals were euthanized and perfused 3 hours later, and brain collected for histological analysis. Detection and quantitative analysis mainly focused on the dentate gyrus (DG) including hilus (H), molecular layer (ML), granular cell layer (GCL), and polymorphic layer (PL). Quantitative analysis was performed on square region highlighted in blue using BZ-X series all in one fluorescence microscope under 2-photon imaging software. Figure generated with BioRender. **B**) Mitophagy in granule cell regions of cerebellum (**B1, B5**), layers 1-3 of the Brodmann area 11 pre-frontal cortex (**B2, B6**), and molecular/granule cell/polymorphic/hilus of DG (**B3, B7**) in male saline injected mito-QC mice. Shown are representative coronal images, with and without Hoechst signal. **B4**) Magnified region from orange box of **B3**) denoting mitophagy puncta by arrows. **B8**) Quantification of mitophagy events per region (340 µm x 260 µm), n=5 each group, 3 month old males. *In assessing the differences among the three data sets, a two-way ANOVA was conducted, complemented by Brown-Forsythe and Bartlet’’s tests for equal variances, and Tuke’’s multiple comparison test for post-hoc analysis. Data passed Kolmorogorov-Smirnov test for normality and no outliers were excluded*. All analysis performed simultaneously using identical brightness, contrast, and exposure. Scale bar=20 µm.

### Immunostaining

Brains were collected, fixed with 4% paraformaldehyde, dehydrated with 30% sucrose, and frozen in OCT before sectioning. We used 10 µm sections for immunofluorescence staining. The following primary antibodies were used: *O*-GlcNAc (1:1000 mouse, Abcam ab2739, RRID:AB_303264), NeuN (1:600 mouse, Millipore MAB377, RRID:AB_2298772), IBA1 (1:600 rabbit, Fujifilm Wako 019–19741, RRID:AB_839504), TMEM119 (1:600 rabbit, Cell Signaling, Cat. No. #90840S), GFAP (1:1000 rabbit, Dako, GA52461-2, RRID:AB_2811722), S100B (1:600 mouse, Proteintech, 66616-1-lg, RRID:AB_2881976), and LAMP1 (1:1000 rat, Abcam, ab25245, RRID:AB_449893). Secondary antibodies were: Alexa Fluor 488 (1:1000 anti-rabbit, Invitrogen: A-10680, RRID:AB_2534062), Alexa Fluor 647 (1:1000 anti-mouse, Invitrogen: A-21235, RRID:AB_2535804), Alexa Fluor 647 (1:1000 anti-rabbit, Invitrogen: A-21245, RRID:AB_141775), and Alexa Fluor 647 (1:1000 anti-rat, Invitrogen: A-21247, RRID:AB_141778). Sections were mounted on Fischer SuperFrost Plus microscope slides. Hoechst 33258 (RRID:AB_2651133) nuclear staining was performed.

### Imaging analyses

Imaging was performed using the Keyence BZ-X810 x40 lenses, centered and localized to the identical region of dentate gyrus for each sample, pictured using sectioning software function, and with identical exposure time for each channel. Analysis was performed using Fiji ImageJ, quantified using measurement and co-localization software tools, with negative control image background values subtracted. Total mitophagy events (punctate red dots) that were ≥0.5 µm diameter were counted. Images were visualized and quantified with an algorithm using consistent brightness and contrast parameters.

### Statistics

GraphPad Prism software (GraphPad Software, Inc., La Jolla, CA, USA) was used for statistical analysis. Two-tailed unpaired t-tests were used to compare the means of two independent groups, calculating the probability that any difference is due to chance without assuming directionality of difference. This analysis was and then followed by an F-test that compares variances by dividing the variance of one data set by the variance of another, testing the null hypothesis that both data sets have the same variance. ANOVA was used to determine if there were any statistically significant differences between the means of <u>three or more</u> independent (unrelated) groups. Brown-Forsythe and Bartlett’s tests were then used to assess the assumption of equal variances among groups analyzed with ANOVA, while Tukey’s multiple comparison test was employed for post-hoc analysis to determine which specific group means were significantly different from each other after finding a significant F-statistic.

## Results

### Mitophagy in dentate gyrus (DG)

To assess the level of mitophagy, the number of red puncta in the cerebellum (**Fig. 1B1, 5**), pre-frontal cortex (**Fig. 1B2, 6**), and the area of dentate gyrus (DG) (**Fig. 1B3, 7**) as depicted in **Fig. 1A2**, were determined in the mito-QC mice. **Fig. 1B4** shows a representative magnified view (from the box area of **Fig. 1B3**) containing the red puncta adjacent to individual cells. We found that the number of red puncta, representative of mitophagy, was highest in the dentate gyrus compared to the cerebellum granular cell layer or the prefrontal cortex in the area shown (**Fig. 1B8**). As expected, the control non-transgenic wild type C57BL/6 did not exhibit red puncta (**Sfig. 1A1-2)**. Consistent with previous reports on the mitophagy of mito-QC mice in the brain ^29^, the mCherry signal co-localized with lysosomal-associated membrane protein 1 (LAMP1) (**Sfig. 1B**). Also agreeing with previous studies, mitophagy occurs in cerebellum Purkinje cells (**Sfig. 1C**).

### Elevation of *O*-GlcNAcylation decreases mitophagy in DG

We next injected (i.p.) male and female mito-QC mice with either saline or the *O*-GlcNAcase inhibitor thiamet G (TG) and collected brains 3 hours later. Using immunohistochemistry, we found that the brains of the TG injected group exhibited the expected increase in protein *O*-GlcNAcylation when compared to saline injected animals (**Fig. 2A-C**). In addition, neurons within the DG granule cell layer (GCL) have the highest *O*-GlcNAc+ staining after TG, in both male and female mice (**Fig. 2A2, 4, & 2B2, 4**). A small proportion of *O*-GlcNAc+ cells are in the molecular layer, which exist in both saline and TG animals, shown in **Fig. 2B3-4**. Interestingly, there was also a significant decrease in the total mitophagy events within *O*-GlcNAc+ cells over the region (**Sfig. 1D**).

**Fig. 2.**
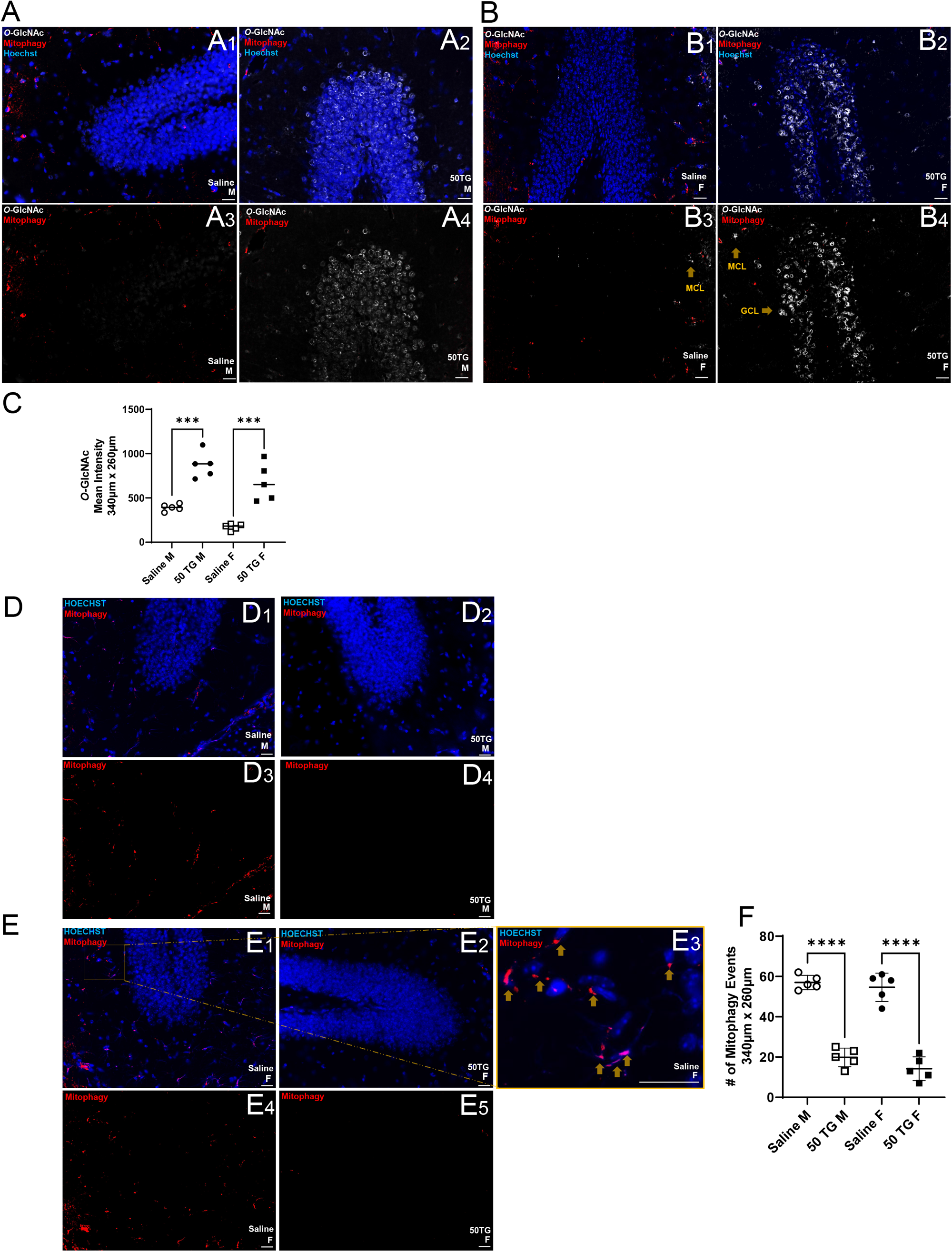
Thiamet G (TG) increased *O*-GlcNAcylation and decreased mitophagy. **A1-4**) Representative images of dentate gyrus from saline and 50 mg/kg TG treated male mito-QC mice stained with an *O*-GlcNAc antibody (white) (n=5). **A1-2, B1-2** are with Hoechst, and **A3-4, B3-4** are without Hoechst. **B1-4**) Representative images of dentate gyrus region from saline and 50 mg/kg TG treated female mito-QC mice (n=5) stained with an O-GlcNAc antibody. **C)** Quantified fluorescence intensity of *O*-GlcNAc immunostaining in mito-QC mice with saline and TG in male and female mice. **D1-4**) Representative images of dentate gyrus region from saline and 50 mg/kg TG treated male mito-QC mice (n=5). **D1-2, E1-3** are with Hoechst, **D3-4**, and **E4-5** are without. **E3)** Magnified region from **E2** orange box, with mitophagy marked by yellow arrow. **F)** Quantification of mitophagic events in male and female mice. All immunohistochemistry analyses were performed simultaneously and imaged in an area of 340 µm x 260 µm, using the same brightness, contrast, and exposure settings over a period of two days. *Two-way ANOVA, Brown-Forsythe, and Bartlett’s test employed for C & F. Data passed Kolmorogorov-Smirnov test for normality and no outliers were excluded. ***p< 0*.*001, ****p< 0*.*0001*. Scale bar=20 µm.

We additionally observed total mitophagy changes across all cell types in DG (**Fig. 2D-E**). We found significantly decreased total mitophagy events after TG in both sexes (**Fig. 2D2, 4 & 2E2, 5**) when compared to respective saline controls (**Fig. 2D1, 3 & Fig. 2E1, 4**). **Fig. 2E3** shows the magnified view from the box in **Fig. 2E1**, with mitophagic puncta marked by arrows. We also observed the majority of mitophagy events are within the molecular layer (**Fig. 2D3, 2E4)**. We quantified the total number of mitophagy events over the image field for both sexes, comparing saline versus TG (**Fig. 2F**).

### TG treatment decreases GFAP+ signal and mitophagy events in GFAP+ cells in DG

Using glial fibrillary acidic protein (GFAP) immunohistochemistry, which detects an intermediate filament protein unique to astrocytes that is also upregulated during neuroinflammation ^31^, we observed that the GFAP level in the DG decreases significantly after 50 mg/kg TG (**Fig. 3A3, 6 & Fig. 3B2, 4**) when compared to after saline in both male and female mice (**Fig. 3A1, 4 & Fig. 3B1, 3**). We quantified GFAP intensity in **Fig. 3C1**. We found that total numbers of GFAP+ cells in the image area were decreased after TG in both male and female mice, with the effect more pronounced in female mice (**Fig. 3C2**).

**Fig. 3.**
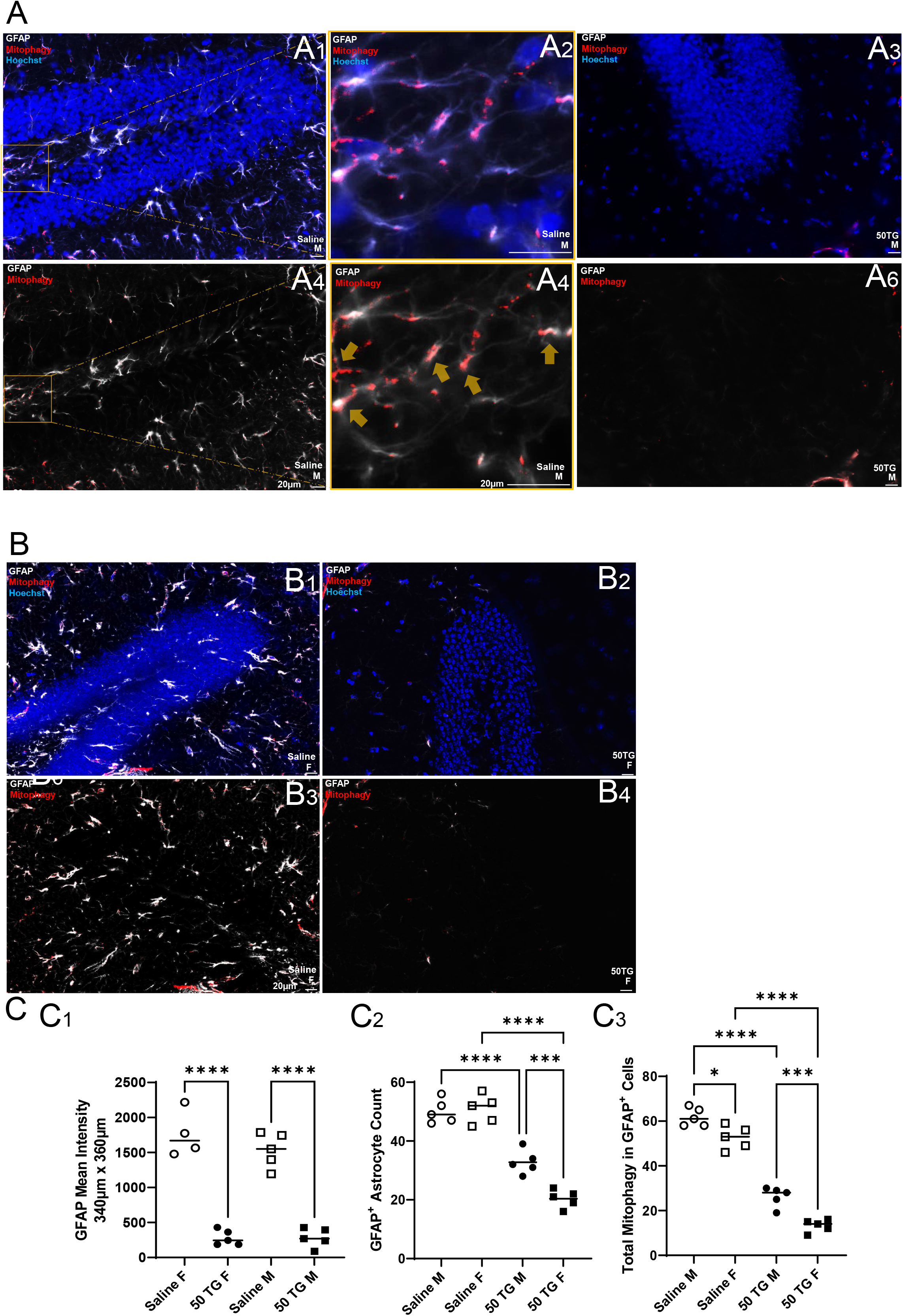
Thiamet G (TG) decreased GFAP+ cell number and mitophagy in GFAP+ cells. **A1-6, B1-4**) Representative mito-QC histological images of DG in male or female saline or 50 mg/kg TG (n=5) treated mice, stained with GFAP in white. **A1-3, B1-2** are with Hoechst, and **A4-6, B3-4** are without Hoechst. **A2, 5**) Representative magnified GFAP+ processes from orange box of **A4**, showing colocalized with mitophagic puncta, marked by arrows. **C1**) Quantified fluorescence GFAP intensity in indicated groups of mice. **C2**) Quantification of the number of GFAP+ cells in indicated groups of mice. **C3**) Quantification of total mitophagy colocalized with GFAP. All immunohistochemistry analyses were performed simultaneously and imaged in an area of 340 µm x 260 µm, using the same brightness, contrast, and exposure settings over a period of two days. *Two-way ANOVA, Brown-Forsythe, and Bartlett’s test employed for C1-3. Data passed Kolmorogorov-Smirnov test for normality and no outliers were excluded*. Scale bar=20 µm.

Additionally, we found that the total number of mitophagy events in GFAP+ cells, as shown by arrows in the representative magnified image (**Fig. 3A5**), decreased after TG (**Fig. 3C3**). There was also less total mitophagy events in GFAP+ cells in female mice compared to male mice, both with saline and TG injection, with more significant difference post-TG (60% down versus 10% down in saline group) (**Fig. 3C3**).

### TG treatment decreases S100B+ signal and mitophagy events in S100B+ cells in DG

We next examined the levels of S100 calcium-binding protein B (S100B) (**Fig. 4A-B**), an astrocyte marker and sensitive inflammation sensor that rises before alterations in intracerebral pressure or during the acute brain injury (22, 23). As with GFAP, S100B levels are lower after TG injection in both male and female mice (**Fig. 4A3, 6 & 4B2, 4**), when compared to saline (**Fig. 4A1, 4 & 4B1, 3**). We quantified S100B immunostaining intensity in **Fig. 4C1**. Additionally, we found the number of S100B+ astrocyte cells decreased after TG injection in both male and female mice (**Fig. 4C2**). We also found that mitophagy events in S100B+ cells, visualized by arrows in the representative magnified images (**Fig. 4A2, 5**), were decreased in male and female mice (**Fig. 4C3**).

**Fig. 4.**
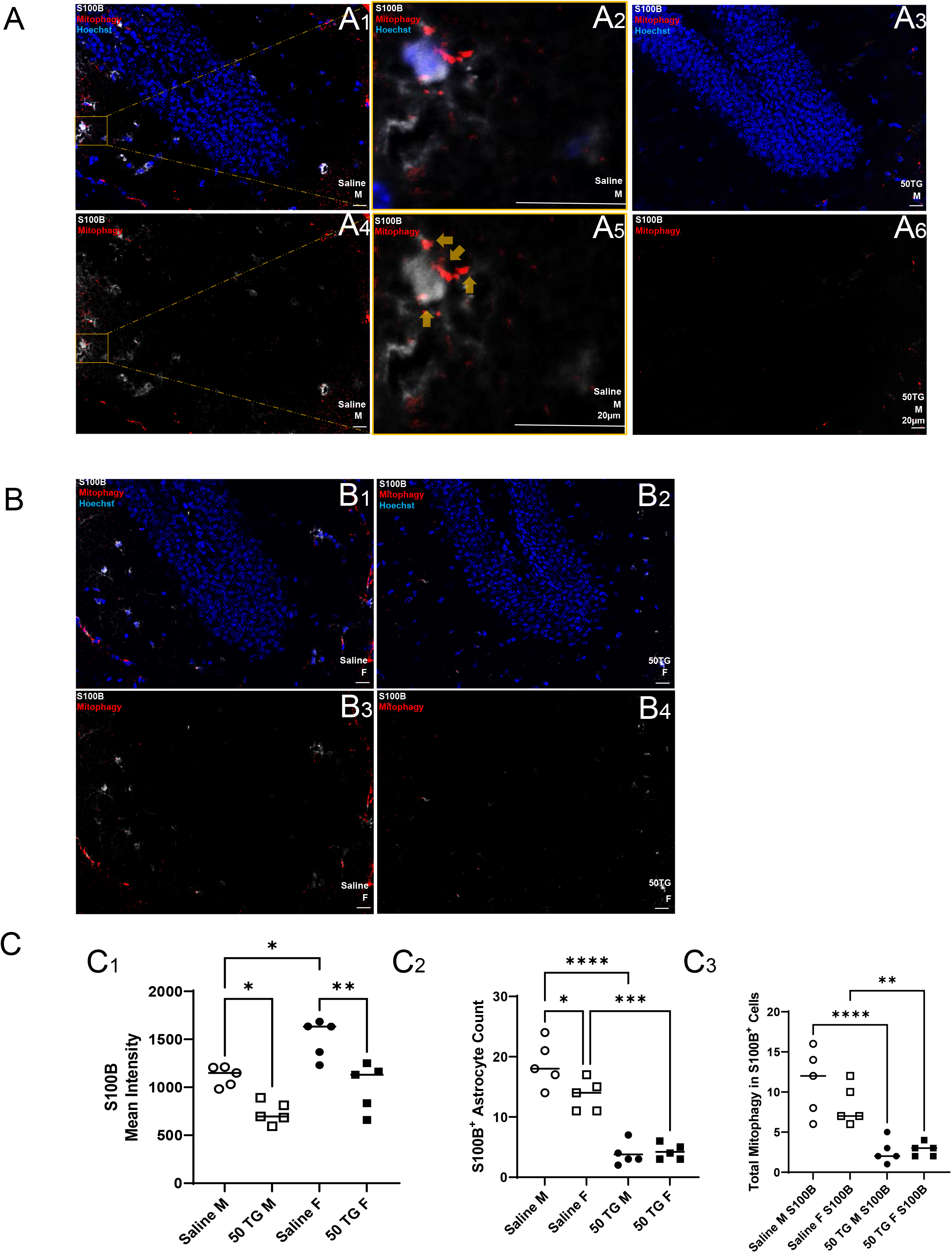
Thiamet G (TG) decreased S100B+ cell number and mitophagy in S100B+ cells. **A1-6, B1-4**) Representative mito-QC histological images of saline or 50 mg/kg TG treated male and female mice (n=5) stained with S100B in white. **A1-3, B1-2** are with Hoechst, and **A4-6, B3-4** are without Hoechst. **A2**,**5**) Representative magnified images from the boxed areas in **A1**,**4**, showing S100B+ processes and soma colocalized with mitophagic puncta, marked by arrows. **C**) Quantified S100B fluorescence intensity **C1**), number of S100B+ cells **C2**), and total mitophagy colocalized with S100B+ cells **C3**), in indicated groups of mice. *Two-way ANOVA, Brown-Forsythe, and Bartlett’s test employed C1-3. Data passed Kolmorogorov-Smirnov test for normality and no outliers were excluded*. Scale bar=20 µm.

### TG treatment decreases IBA+ signal while increases total mitophagy events in IBA+ cells in DG in male mice

We then investigated whether microglia are affected by acute TG injection. To accomplish this, we performed immunohistochemistry with an antibody against ionized calcium-binding adapter molecule 1 (IBA1), a known marker for activated / inflammatory microglia and upregulated during nerve injury, ischemia, exposure to IFN-γ, IL-1β, and T-cell activation (24) (**Fig. 5**). As with GFAP and S100B, IBA+ staining was significantly decreased in the DG of TG treated mice (**Fig. 5A3, 6 & B2, 4)** compared to saline control (**Fig. 5A3, 6 & B2, 4**), which we quantified in **Fig. 5C1**. We also noted that mitophagic puncta were frequently adjacent to, and rarely colocalize with, IBA1+ processes, marked by arrows in the representative magnified image (**Fig. 5A5**, from boxed area of **Fig. 5A4**). Additionally, we detected a significant decrease in the number of IBA1+ microglial cells, more in female mice (**Fig. 5C2**). The total IBA1+ cells with mitophagy events after TG only increased in male mice (**Fig. 5C3**).

**Fig. 5.**
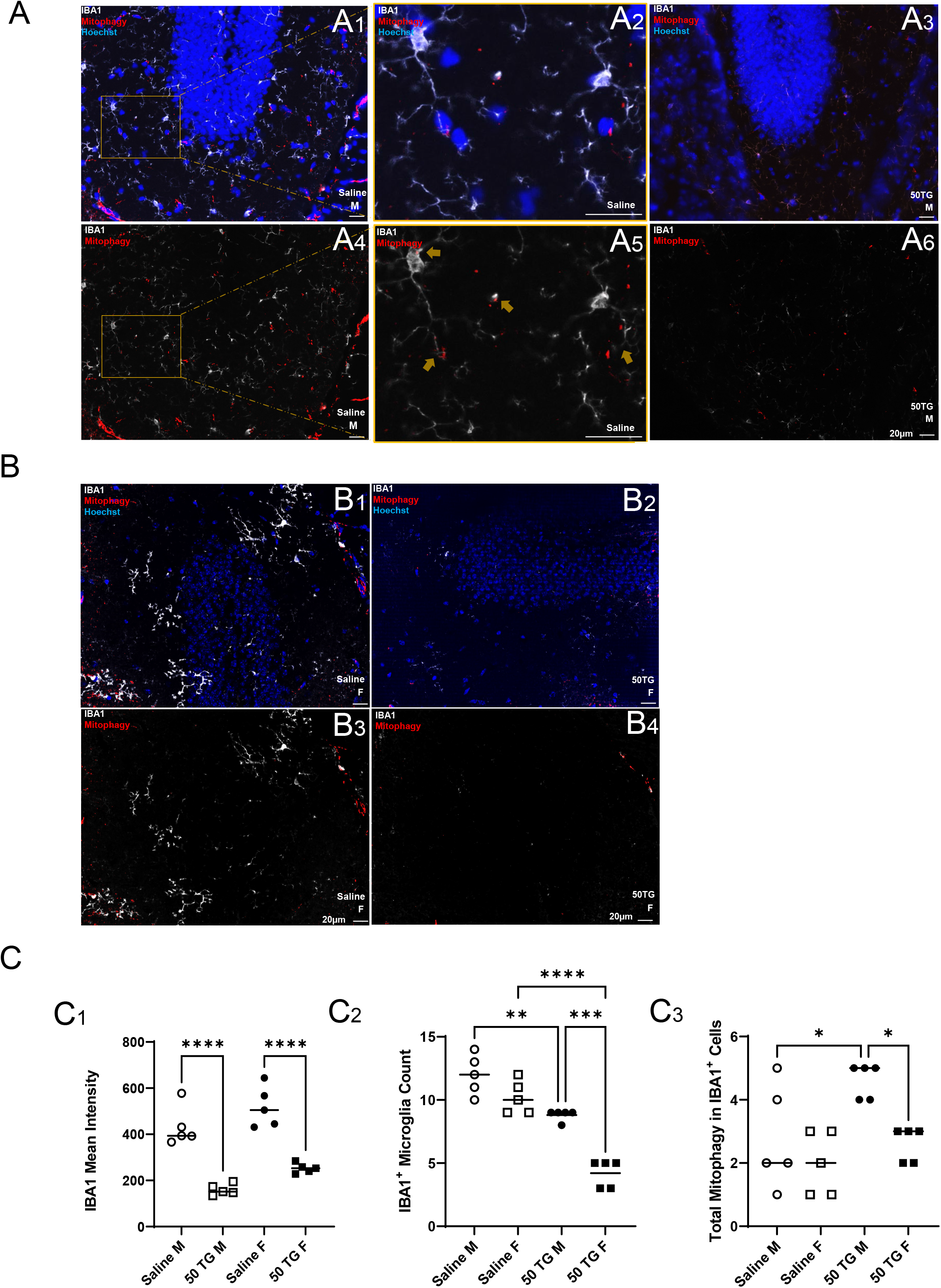
Thiamet G (TG) decreased IBA1+ cell number. **A1-6, B1-4)** Representative mito-QC histological images of male and female saline or 50 mg/kg TG (n=5) treated mice stained with IBA1 in white. **A1-3, B1-2** are with Hoechst, while **A4-6 & B3-4** are without Hoechst. **A2, 5** are magnified images from the boxed areas in A1,4, showing IBA1+ processes adjacent to and colocalized with mitophagic puncta, marked by arrows. **C1**) Quantified IBA1 fluorescence intensity, **C2**) Number of IBA1+ cells, and **C3**) Total numbers of mitophagy event in IBA1+ cells, in indicated groups of mice. *Two-way ANOVA, Brown-Forsythe, and Bartlett’s test also employed in C1-3. Data passed Kolmorogorov-Smirnov test for normality and no outliers were excluded*. Scale bar=20 µm.

### TG treatment increases TMEM119+ signal

We next examined immunostaining of transmembrane protein 119 (TMEM119) (**Fig. 6**), which is frequently used to mark homeostatic or alternatively activated/anti-inflammatory microglia that have been shown to produce neurotrophic factors benefiting regeneration and clearance of debris inhibiting tissue repair (25, 26). In contrast to IBA1, TMEM119 immunostaining intensity was significantly increased after TG (**Fig. 6A2, 4 & B3, 6**) when compared to saline controls (**Fig. 6A1, 3 & B1, 4**), which we quantified in **Fig. 6C1**. As with IBA1 in saline treated animals, TMEM119+ processes were also often found adjacent to mitophagic puncta (**Fig. 6B2, 5**) magnified from boxed area in **Fig. 6B4, 6**).

**Fig. 6.**
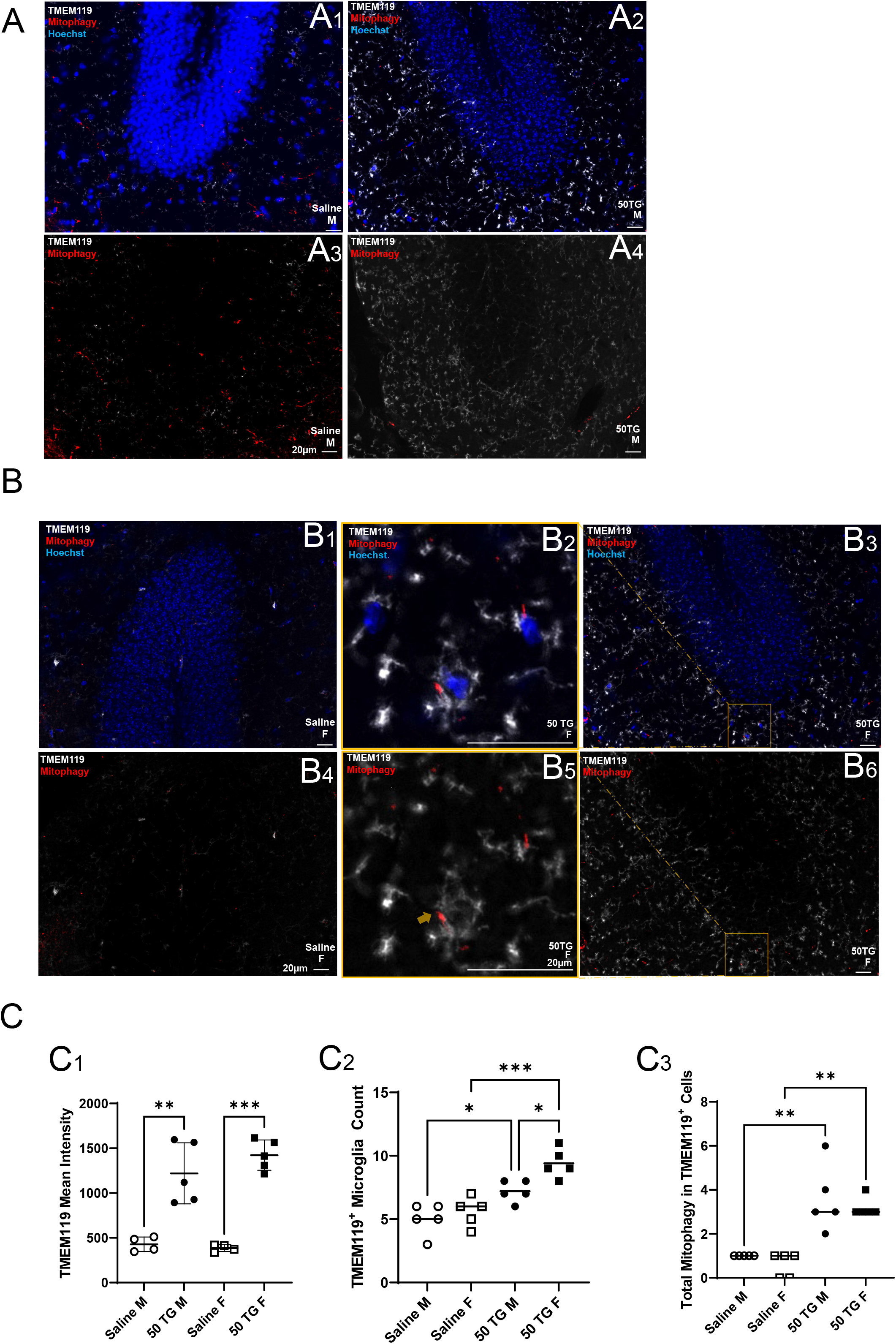
Thiamet G (TG) increased TMEM119+ cell number and mitophagy in TMEM119+ cells. **A1-4, B1-6**) Representative histological images of saline or 50 mg/kg TG (n=5) treated male and female mito-QC mice, stained with TMEM119 in white. **A1-2 & B1-3** are with Hoechst, while **A3-4 & B4-6** are without Hoechst. **B2, 5**) Magnified images from orange box of **B3, 6**, showing TMEM119+ processes and nearby mitophagic puncta, marked by arrows. **C1**) Quantified TMEM119 fluorescence intensity, **C2**) Number of TMEM119+ cells, and **C3**) Total numbers of mitophagy events in TMEM119+ cells, in indicated groups of mice. *Two-way ANOVA, Brown-Forsythe, and Bartlett’s test also employed in C1-3. Data passed Kolmorogorov-Smirnov test for normality and no outliers were excluded*. Scale bar=20 µm.

Concurrent with elevated TMEM119 intensity, the number of TMEM119+ microglial cells were also increased, with females exhibiting both a higher number of TMEM119+ cells after TG and a more significant increase (82% increase in females versus 44% in males) (**Fig. 6C2**). Furthermore, the total number of mitophagy in TMEM119+ cells in both sexes was also increased (**Fig. 6C3**).

### TG treatment does not change NeuN+ intensity in DG

To assess the impact of TG on neuronal markers in the DG, we stained for neuronal nuclear antigen (NeuN), which binds to a protein primarily found in the nucleus of neurons (**Fig. 7**). We found no significant difference between NeuN+ intensity or NeuN+ cell number in saline and TG treatment groups of either male or female mice (**Fig. 7A1-4, B1-6**), which we quantified in **Fig. 7C1-2**. Mitophagic puncta were adjacent to NeuN+ cells in the ML of the DG, denoted by arrows in the representative magnified image (**Fig. 7B2, 5**). The colocalization of NeuN with mitophagy was rare (**Sfig. 1E**). However, there was a decrease in the total number of mitophagy events in NeuN+ cells (**Fig. 7C3**).

**Fig. 7.**
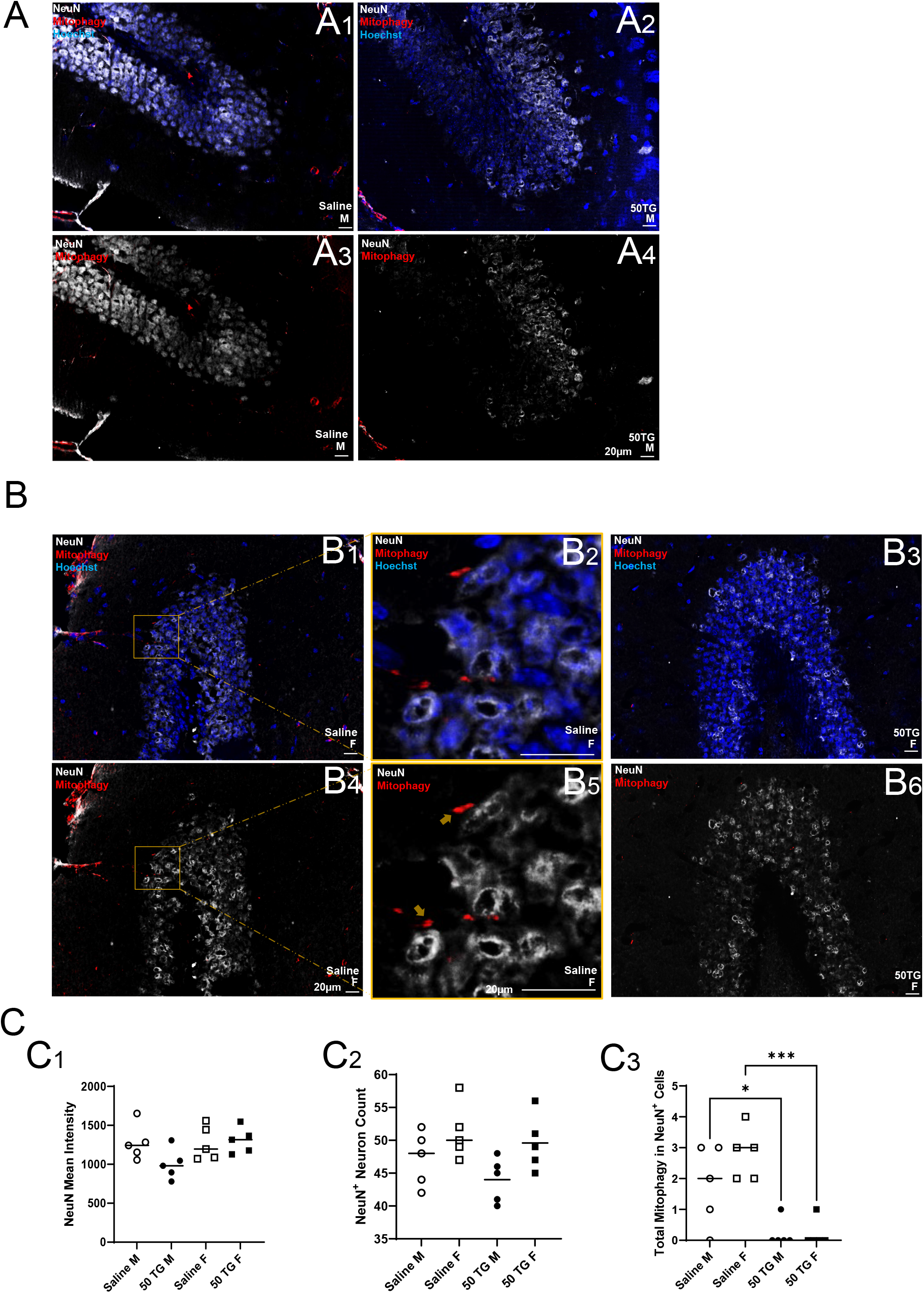
NeuN+ immunostaining intensity is unchanged by thiamet G (TG). **A1-4, B1-6)** Representative histological images of male and female mito-QC saline or 50 mg/kg TG treated mice (n=5), stained with NeuN in white. **A1-2 & B1-3** are with Hoechst, while **A3-4 & B4-6** are without Hoechst. **B2, 5**) Magnified image from orange box of **B1, 4**) showing NeuN+ soma shown adjacent to mitophagic puncta, marked by arrows. **C1)** Quantified NeuN fluorescence intensity, **C2**) Number of NeuN+ cells, and **C3**) Total mitophagy events colocalizing with NeuN+ signal, in indicated groups of mice. *Two-way ANOVA, Brown-Forsythe, and Bartlett’s test employed for C1-3. Data passed Kolmorogorov-Smirnov test for normality and no outliers were excluded*. Scale bar=20 µm.

Considering the total mitophagy events in male and female mice are ∼60 in saline treated mice (**Fig. 2F**), we conclude, that under saline injections, neuronal and microglial mitophagy is less prominent than mitophagy in GFAP+ astrocytes (**Fig. 8A1**). After TG injections, there are an average of approximately 20 mitophagy events in the dentate gyrus region (**Fig. 2D6**), and we find that GFAP+ astrocytes still make up the majority of those events (**Fig. 8A2**).

**Fig. 8.**
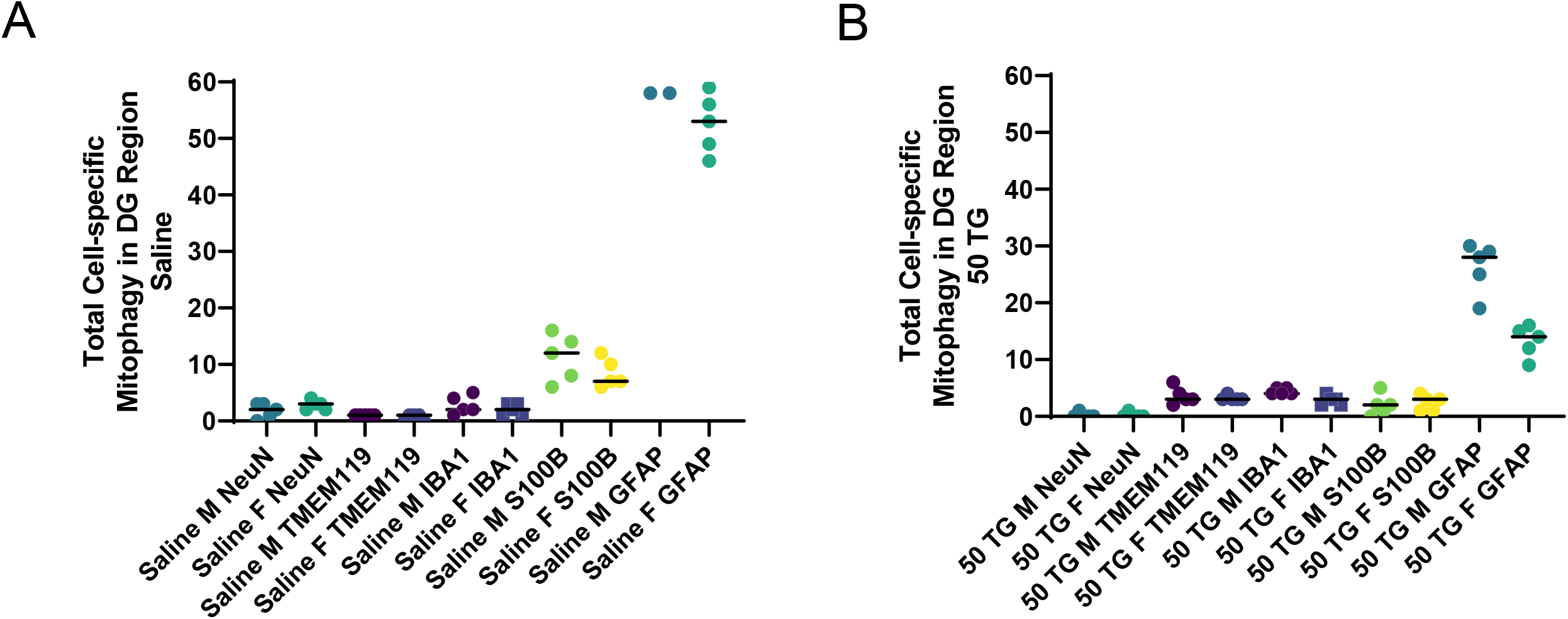
Astrocytes contain the majority of total mitophagy events in DG. **A**) Comparison of total mitophagy event colocalizing with specific cell markers under saline condition. **B**) Comparison of total mitophagy event colocalizing with specific cell markers under 50 TG condition. *Data passed Kolmorogorov-Smirnov test for normality and no outliers were excluded*. Scale bar=20 µm.

### TG treatment in 6- and 26-month-old mice decreases GFAP+ immunostaining intensity

Aging induces a myriad of molecular and cellular transformations, potentially impacting the response to elevation of *O*-GlcNAcylation within the brain. Consequently, we investigated the patterns of GFAP+ immunostaining within the DG of 6 and 26 month old mice with saline and TG injection to compare to our findings in younger mice. We found that TG significantly decreased GFAP+ immunostaining intensity in both 6 month (**Fig. 9A1-4**) and 26 month old mice (**Fig. 9B1-4**), quantified in **Fig. 9C**. We also observed that in 26-month-old mice, the decrease of GFAP immunostaining intensity after TG is larger than in younger mice, driven by higher basal level of GFAP in 26-month-old mice (**Fig. 9C**). *O*-GlcNAcylation immunostaining (data not shown) also confirmed that TG elevated *O*-GlcNAcylation in both 6- and 26-month-old mice.

**Fig. 9.**
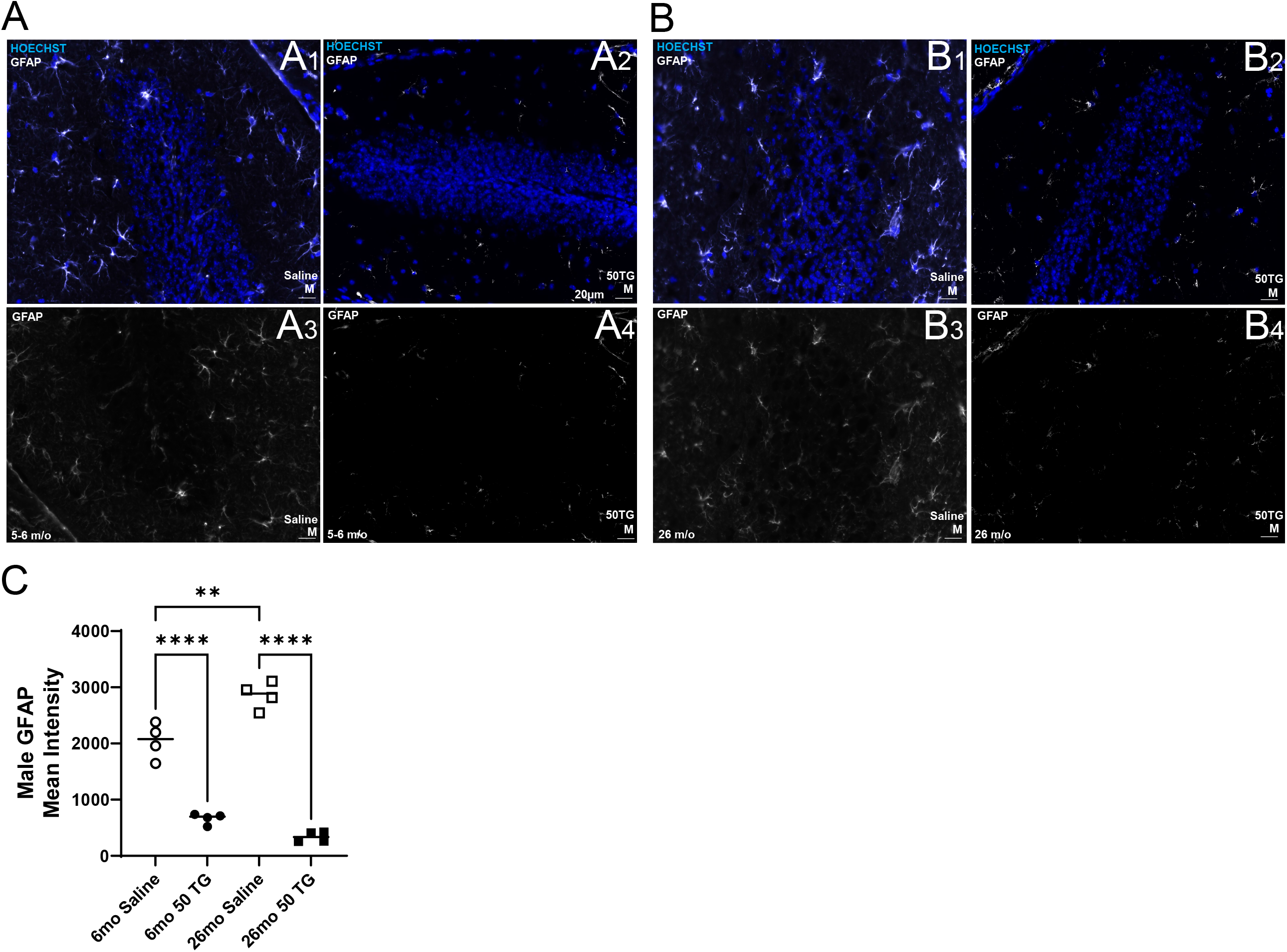
Thiamet G (TG) decreased GFAP and increased *O*-GlcNAc in 6 and 26 month old male mice. **A1-4, B1-4**) Representative histological images of DG of saline or 50 mg/kg TG (n=4) treated C57BL6 6 mo and 24 mo male mice, stained with GFAP in white. **A1-2 & B1-2** are with Hoechst, while **A3-4 & B3-4** are without Hoechst. **C**) Quantified GFAP fluorescence intensity in DG in indicated groups of mice. Two-way *ANOVA, Brown-Forsythe, and Bartlett’s test employed in C. O-GlcNAc immunostaining not shown. Data passed Kolmorogorov-Smirnov test for normality and no outliers were excluded*. Scale bar=20 µm.

## Discussion

Our study identified basal mitophagy in astrocytes, microglia, and neurons in the DG in 3-month-old male and female mice. Furthermore, we demonstrate *in vivo* an early response to OGA inhibition is a profound decrease in mitophagy in the DG in both sexes. This decrease in mitophagy also corresponds with significant changes in the glial cell composition. Elevation of *O*-GlcNAcylation decreased astrocyte markers and mitophagy events in cells expressing these markers in both male and female mice. Elevation of *O*-GlcNAcylation also resulted in specific changes in microglia, decreasing IBA1 level and IBA1+ cell counts, while increasing the level of TMEM119 and TMEM119+ cell count (**Fig. 10**). These results were more pronounced in females compared to males, a phenomenon potentially attributable to sex-dependent regulation of *O*-GlcNAc recycling, given that the OGT enzyme is located at the X-chromosome ^32^.

**Fig. 10.**
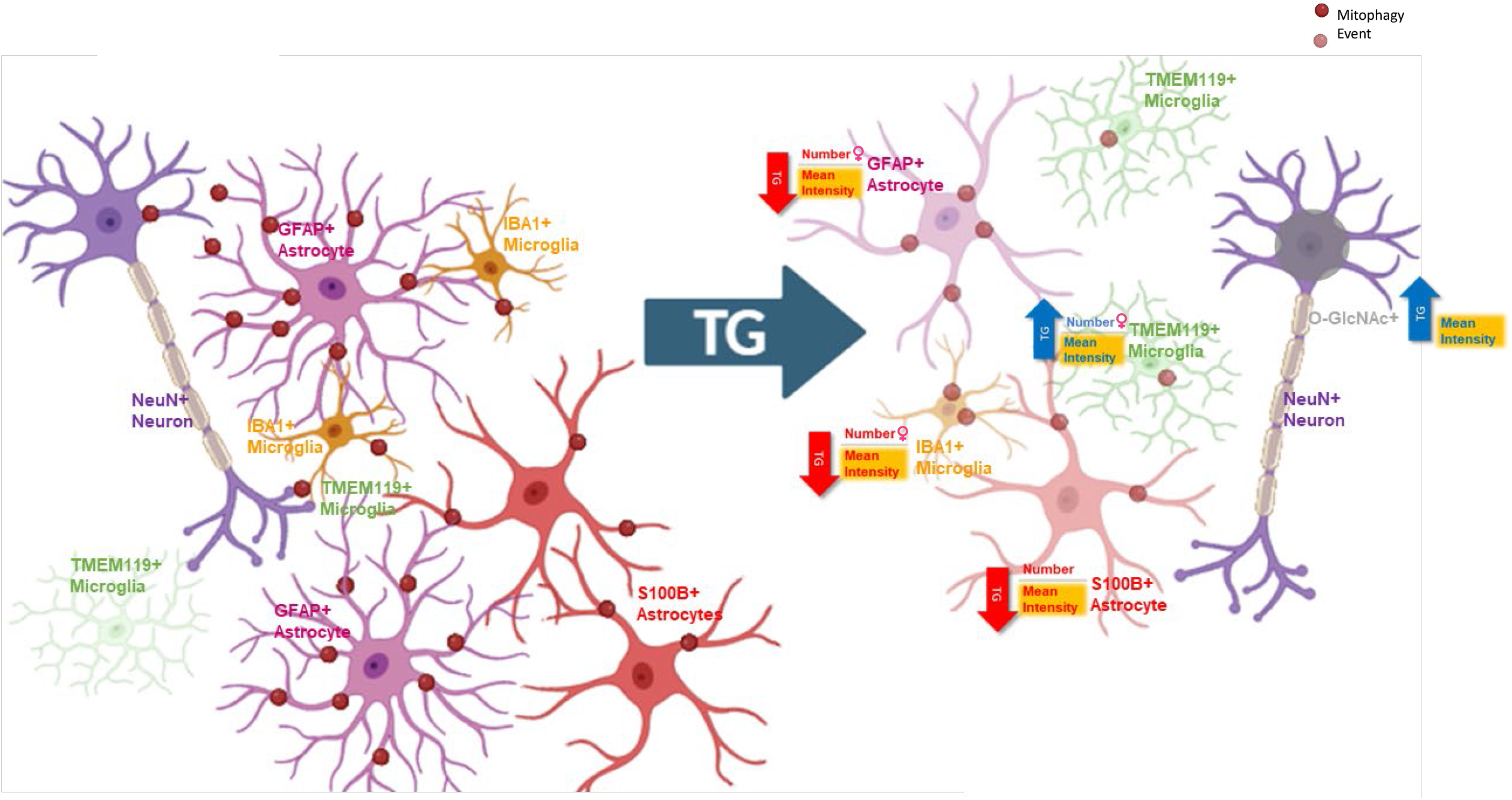
Summary of effects of acute elevation of O-GlcNAcylation by TG on mitophagy and glia homeostasis in the dentate gyrus. This study demonstrates that basal mitophagy occurs mainly in GFAP+ cells, and that acute TG treatment suppresses mitophagy, decreases GFAP+, S100B+, and IBA1+ cell numbers and increases TMEM119+ cell numbers. In addition, there were decreased mitophagy in NeuN+, GFAP+ and S100B+ cells, and no change in TMEM119+ and IBA1+ cells, although percent of TMEM119+ and IBA1+ cells with mitophagy were increased. Female icons indicate more pronounced changes in female mice for numbers of GFAP+, IBA+ and TMEM119+ cells. Figure generated with BioRender.

### The majority of mitophagy events occur In GFAP+ astrocytes in DG

Under physiological conditions in the dentate gyrus (DG), neurons and GFAP+ astrocytes are the major cell types, followed by S100B+ astrocytes, IBA1+ and TMEM119+ microglia. Notably, the majority of mitophagy events occur in GFAP+ astrocytes, exceeding those in microglia and neurons. Even though NeuN primarily stains neuronal cell bodies, excluding axons, the majority of mitophagy events still occur in GFAP+ cells, as evidenced by the quantifications in **Fig. 2F** and numbers of mitophagy event colocalized with different cell markers in **Fig. 8A-B**.

Given the energy dependence of neuronal activities and the largely post-mitotic nature of neurons, the efficient removal of dysfunctional mitochondria is vital for their health. This leads to the possibility that neuronal mitochondria are degraded by astrocytes and microglia. This observation is consistent with prior studies that astrocytic mitophagy can be important for neuronal mitochondrial quality control, through a process known as transmitophagy, where damaged mitochondria from neurons are transferred to astrocytes for degradation ^31^. Additionally, transmitophagy might also occur in microglia, analogous to macrophages preserving cardiomyocyte mitochondrial integrity during cardiac ischemia through mitochondria sharing via vesicles ^33^. Studies in *C. elegans* have shown that under stress, neurons can expel large vesicles, known as exophers, containing organelles like mitochondria, an effect crucial for waste elimination and potentially beneficial in treating neurodegenerative diseases ^34-37^, underscoring a need for further research into these mechanisms.

### Elevation of *O*-GlcNAcylation changes the glial cell composition

One surprising and interesting finding from our study is that 3-hour elevation of *O*-GlcNAcylation significantly changes the glia cell composition. Specifically, elevation of *O*-GlcNAcylation significantly decreased the numbers of GFAP+ and S100B+ astrocytes, and IBA1+ microglia, while increasing the numbers of TMEM119+ microglia. The mechanisms behind this significant change are currently unclear. One possibility is that elevation of *O*-GlcNAcylation changes the expression of these glial cell markers. It is known that elevated protein *O*-GlcNAcylation can change the transcription profile ^38^. Prior studies demonstrate that increasing *O*-GlcNAc levels modify Sp1 transcription factor and increase the transcription of glycerol-3-phosphate acyltransferase-1 (GPAT1) ^38^. Future studies using scRNAseq will be crucial to determine the cell specific effect of elevation of *O*-GlcNAcylation on the glial composition, and to determine how elevation of *O*-GlcNAcylation affects phagocytic and neuronal supportive capacity, as well as neuroinflammation response in neurodegenerative diseases including AD.

### Elevation of *O*-GlcNAcylation decreases mitophagy and this is largely mediated by a decrease of GFAP+ cells

Another significant finding from our study is that elevation of *O*-GlcNAcylation *in vivo* immediately induced a dramatic decrease of mitophagy in the DG. Considering that elevation of *O*-GlcNAcylation significantly decreased the numbers of GFAP+, and S100B+ cells, this decrease of mitophagy may be due to a significant change of transcriptional landscape in these cells resulting in downregulation of mitophagy in these cells. Additionally, even though NeuN+ cell numbers were unchanged by elevation of *O*-GlcNAcylation, mitophagy in NeuN+ cells also decreased. In contrast, although IBA1+ cell number was decreased, mitophagy events in male IBA1+ cells were increased, while TMEM119+ cell number and mitophagy events in TMEM119+ cells were also increased. While the potential mechanism is currently unclear, this may be due to the different transcriptional and post-transcriptional regulation of genes involved in mitophagy in different cells.

### Elevation of *O*-GlcNAcylation also decreases GFAP immunoreactivity in aged mice

We further demonstrated that significant immediate downregulation of GFAP by elevation of *O*-GlcNAcylation also occurred in non-mito-QC transgenic mice at 6 and 26 months of age. There was significantly elevated GFAP+ signal in 26 months old mice compared to 6 months of mice. Nonetheless, in response to elevation of *O*-GlcNAcylation, detected by immunostaining (data not shown), there was significantly decreased GFAP+ signal. Whether the acute changes in GFAP by elevated *O*-GlcNAcylation is pro or anti-inflammatory remains to be tested, which is crucial for fully understanding their roles in aging and neurodegenerative diseases.

### Limitations of this study

There are several limitations of our study that could inform future research directions. First, the mito-QC reporter used in our experiments, while effective, is less sensitive compared to mt-Keima ^39^, potentially affecting the detection sensitivity of mitophagy events. Additionally, our focus was exclusively on the dentate gyrus (DG), other brain regions may not have the same cell type distribution of mitophagy events and the response to elevation of *O*-GlcNAcylation. Also, since we observed that the majority of mitophagy events were in GFAP+ cells, in this study we did not investigate mitophagy in other cell types in the brain, including oligodendrocytes, endothelial cells, or neuronal axons.

Furthermore, our study was conducted with injecting TG at a single time of day and harvesting the brain 3 hours post-injection. Future studies may provide further insights of whether TG effect is time-of-day dependent. In the case of 26-month-old mice, we did not include assessments for IBA, TMEM119, or S100B, nor did we examine female mice, which could have provided further insights into age and sex-related differences in mitophagy.

## Conclusions

In summary, our study demonstrates for the first time, basal level of mitophagy occurs mainly in GFAP+ astrocytes in the DG in both male and female mice, which is dramatically decreased in response to acute treatment with TG and the associated increased protein *O*-GlcNAcylation. This decrease in mitophagy is also concurrent with significant changes of glial cell milieu. Our study provides new insights into the extent of basal mitophagy in different cell types in the brain and the regulation of mitophagy and glia landscape by elevation of *O*-GlcNAcylation. Future studies will determine the mechanisms of the responses to *O*-GlcNAc elevation and the impact of cell specific mitophagy and glial landscapes on mitochondrial function, neuroinflammation, aging and neurodegenerative diseases.

## List of Abbreviations

AD: Alzheimer’s disease
BrdU: Bromodeoxyuridine
COX2: Cyclooxygenase 2
DG: Dentate gyrus
GFAP: Glial Fibrillary Acidic Protein
GFP: Green Fluorescent Protein
GL: granular layer
IBA1: Ionized calcium-binding adapter molecule 1
iNOS: Inducible nitric oxide synthase
LAMP1: Lysosomal-associated membrane protein 1
LC3: Microtubule-associated protein 1A/1B-light chain 3
mCherry-GFP: refers to a fusion protein of mCherry and GFP
Mito-QC mice: Mitochondria Quality Control reporter mice
ML: molecular layer
mo: month
NeuN: Neuronal nuclei
NF-κB: Nuclear factor kappa-light-chain-enhancer of activated B cells
OCT: Optimal Cutting Temperature compound
OGA: *O*-GlcNAcase
*O*-GlcNAc: *O*-linked N-acetylglucosamine
*O*-GlcNAcylation: *O*-linked N-acetylglucosamine modification
OGT: *O*-GlcNAc transferase
PBS: Phosphate-buffered saline
PFA: Paraformaldehyde
PINK1: Phosphatase and tensin homolog induced kinase 1
p62: Sequestosome 1
RRID: Research Resource Identifier
S100B: S100 calcium-binding protein B
TG: thiamet G
TMEM119: Transmembrane protein 119

## Acknowledgment

We thank Dr. Ian Ganley for the mito-QC mice, Drs. Darley-Usmar, Chatham, Young and Zhang lab members for discussion and technical support.

## Author contribution

JK performed experiments. JCC, MEY, VDU, and JZ directed the study. JK and JZ wrote the manuscript. All read/edited/commented/approved the manuscript.

