## supplemental figure for "Impact of *O*-GlcNAcylation elevation on mitophagy and glia in the dentate gyrus"

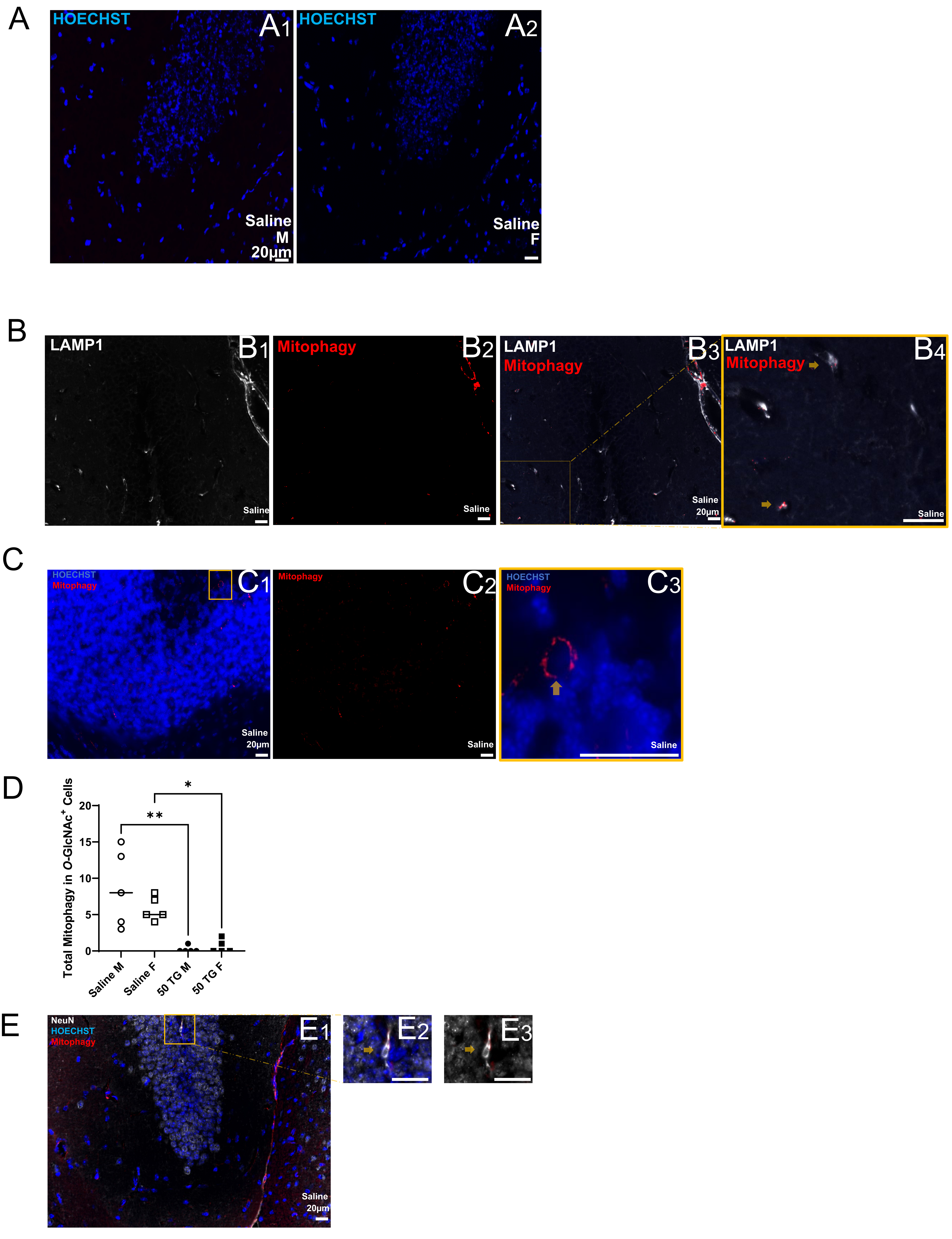
A
HOECHST
A1
Saline
M
20µm
HOECHST
A2
Saline
F
B
LAMP1
B1
Saline
Mitophagy
B2
Saline
LAMP1
Mitophagy
B3
Saline
20µm
LAMP1
Mitophagy
B4
Saline
C
HOECHST
Mitophagy
C1
Saline
20µm
Mitophagy
C2
Saline
HOECHST
Mitophagy
C3
Saline
D
Total Mitophagy in O-GlcNAc+ Cells
20
15
10
5
0
**
*
Saline M
Saline F
50 TG M
50 TG F
E
NeuN
HOECHST
Mitophagy
E1
Saline
20µm
E2
E3

Sfig.1

**Fig.1S. A) Mitophagy mCherry only signals colocalize with lysosomal marker LAMP1. A1-2) Negative control for mitophagy puncta.** Representative in non-transgenic C57BL/6J 3-month-old saline injected mice within the 340 μm x 260 μm region of DG. **A1-2, C1, 3,** & **E1-2** are with Hoechst, while **B1-4, C2** & **E3** are without Hoechst. **B) Red puncta are in the lysosomes.** Lysosomal marker LAMP1 staining was performed in mito-QC mice in the 340 μm x 260 μm DG region of male saline mito-QC mouse. All red puncta are stained with LAMP1 positive signals **B1)** LAMP1 staining (white). **B2)** mCherry red only puncta. **B3)** merged. **B4)** Enlarged from the box in **B3**. **C) Mitophagy detection in Purkinje cells.** Purkinje Cell (340 μm x 260 μm) region adjacent to granular cell layer of male saline mito-QC mouse cerebellum with mitophagic puncta in red, displaying canonical ring structure of mitochondria. **C1)** mCherry red only puncta with nuclear Hoechst staining. **C2)** mCherry red only puncta. **C3)** Enlarged from box in **C1**. **D) Quantification of mitophagy in *O*-GlcNAc+ cells.** Total number of mitophagy events in *O*-GlcNAc+ cells. **E) Representative image of mitophagy colocalization with NeuN. E1** NeuN, Hoechst, and mitophagy signals imaged in 340 μm x 260 μm DG region of saline treated mito-QC male mouse. **E2** shows magnified region from the box in **E1**, and **E3** shows the same region as **E2** without Hoechst for clarity. All analyses were performed simultaneously using identical brightness, contrast, and exposure. Scale bar=20 μm. *Two-way ANOVA, Brown-Forsythe, and Bartlett's test employed in C.* Scale bar=20 μm.
